# Identification and analysis of mobile genetic elements in Gibbon genome

**DOI:** 10.1101/237685

**Authors:** Kamal Rawal, Jaisri Jagannadham, Chahat Kubba, Tanya Sharma

## Abstract

Recent sequencing of genome of northern white-cheeked gibbon (*Nomascus leucogenys*) has provided important insight into fast evolution of gibbons and signatures relevant to gibbon biology. It was revealed that mobile genetic elements (MGE) seems to play major role in gibbon evolution. Here we report that most of the gibbon genome is occupied by the MGEs such as ALUs, MIRs, LINE1, LINE 2, LINE 3, ERVL, ERV-class1, ERV-class II and other DNA elements which include hAT Charlie and TcMar tigger. We provide detailed description and genome wide distribution of all the MGEs present in gibbon genome. Previously, it was reported that gibbon-specific retrotransposon (LAVA) tend to insert into chromosome segregation genes and alter transcription by providing a premature termination site, suggesting a possible molecular mechanism for the genome plasticity of the gibbon lineage. We show that insertion sites of LAVA elements present atypical signals/patterns which are different from typical signals present at insertion sites of Alu elements. This suggests possibility of distinct insertion mechanism used by LAVA elements for their insertions. We also find similarity in signals of LAVA elements insertion sites with atypical signals present at Alus /L1s insertion sites disrupting the genes leading to diseases such as cancer and Duchenne muscular dystrophy. This suggest role of LAVA in premature transcription termination.

## Introduction

*Nomascus leucogenys*, commonly known as white-cheeked gibbon is a member of Hylobatidae family. The gibbons are especially adapted to a tree living lifestyle with a special form of locomotion called brachiation, or hand-over-hand swinging [1]. Gibbon genome was published in 2014 after sequencing of four hominids which includes human, orangutan, gorilla and chimpanzee. They are small arboreal apes closely related to humans and great apes. Comparisons between gibbon chromosomes and those of other primates revealed that gibbons present an extraordinary level of evolutionary chromosomal rearrangements [2].

Mobile genetic elements (MGEs) account for a significant proportion of eukaryotic genomes and play an important role in altered gene expression and disease. It was shown in a comparative study of gibbons with other primates that genomic breakpoints are significantly depleted of genes and breakpoint intervals contained a mixture of repetitive sequences that inserted exclusively into the gibbon genome [2]. Authors also reported that the gibbon genome not only contain several classes of transposable elements as described previously in primate genomes but also contain a novel retrotransposons called as LAVA element [2, 3]. LAVA element is a unique repetitive element that is exclusively present in gibbon genome and is composed of sequences similar to L1, Alu and VA section of SVA mobile elements as shown in figure 1.

**Figure 1:**
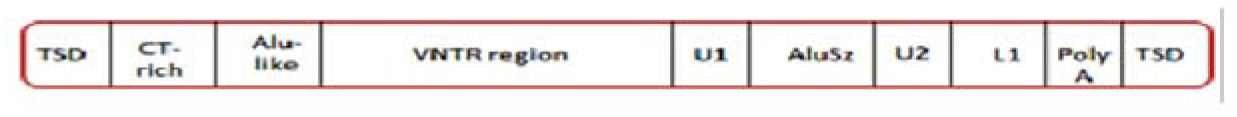
LAVA element structure: LAVA comprises sequences derived from different repetitive elements like L1,Alu-like, and VNTR region.

Earlier studies have also reported that retrotransposons are the most abundant MGE in mammalian genomes which affects wide functional activities such as genome evolution, gene disruption and regulation [4, 5]. The L1 family of LINE elements and Alu elements of SINE are the major class of retroelements believed to impact primate genomes such as gorilla, chimpanzee, and gibbons [6, 7, 8]. These parasitic DNAs have played important roles in the evolution of complex genomes and can also be regarded as the useful tools to study organization, structure and function of eukaryotic genomes. They are generally involved in gene inactivation, gene duplication, transduction and regulation [4, 9]. The recent sequencing of gibbon genome has offered an opportunity to study chromosome wise distribution of these elements.

Till date, very few studies, related to the detailed identification of the transposable elements and their insertion sequences in the gibbon genome have been carried out. The present work involves the identification and classification of different MGEs particularly Alu and LAVA and detailed study of their distribution throughout the genome of *Nomascus leucogenys* using Repeatmasker and ELAN pipeline. ELAN is an efficient computational genomic analysis tool which is responsible for analysing MGEs at different insertion sites whereas Repeatmasker is a program that screens DNA sequences for interspersed repeats [10]. The output of the Repeatmasker program provides detailed classification of the repeats like LTRs, Non-LTRs and other DNA elements. We have also used ELEFINDER (part of ELAN) to find MGEs such as Alu and LAVA in the gibbon genome and also screened the physicochemical properties of insertion sites using DNASCANNER.

## Methods

### Retrieving genome sequences

Gibbon genome was retrieved from NCBI ftp server link ftp://ftp.ncbi.nih.gov/genomes/Nomascusleucogenys/. The chromosomes are made of a total of 2,756,591,777 base pairs of nucleotides.

### Repeat sequence retrieval

Identification of TEs based on their features has enabled the construction of libraries of consensus sequences of various types of TEs known as Repbase. Repbase Update (RU) is a comprehensive database of repetitive element consensus sequences [11].

### Repeatmasker for screening of DNA sequences

Repeatmasker is software that screens DNA sequences for low complexity sequences, repetitive/TEs including small RNA pseudogenes, Alus, LINEs, SINEs, and LTR. Repeatmasker makes use of Repbase libraries which act as standard reference points for the identification of transposable elements in a query sequence. It uses statistically optimal scoring matrices derived from the alignments of DNA transposon fossils to their consensus sequences [10]. However; it does not locate all polymorphic simple repetitive sequences. We used RM-BLAST as search engine in Repeatmasker and tested various options combining speed/sensitivity parameters. Based upon its results, the average length, and the total length occupied by each element was also calculated for each chromosome.

### Whole genome distribution analysis by ELEFINDER

It finds distribution, nature, orientation, genomic location, and site of truncations of mobile genetic elements [8, 12]. ELEFINDER not only identify repeats but also extracts flanking sequence at each MGE site. We used ELAN to find out the insertion sites of various MGEs in the gibbon genome. It is a Perl based system which uses chromosome number, genome file name, and element file name as an input. The results files are generated after performing BLASTN and use of parsers written in Perl. The output files comprise the list of MGEs categorized into full length (intact copies), 5′ truncated, 3′ truncated and both side truncated examples. It also provides upstream sequences for analysis by DNA SCANNER tool.

## DNA SCANNER

The tool DNA SCANNER scans genomic DNA for a variety of properties which includes thermodynamic properties, physicochemical properties, interactions between protein and sequence based features like the AT density [8, 13, 14]. It is also responsible for analysing the insertion sites of the repeats that are known and the detects the presence of different signals. DNA sequence analysis method is basically limited to homology searches such as BLAST, finding the patterns in the sequence that are sequence based and the percentages of GC and AT. It offers another pathway for DNA sequence analysis and has wide pertinence in the analysis of DNA sequences which further allow the analysis of promoters, genic and intergenic regions, insertion sites of retroelements, intron-exon boundaries and clustering sequences based upon their properties [8]. The program evaluates number of properties for a given DNA sequence in a sliding window to generate sequence based profiles as explained below.

A. Structural signals: DNA bendability

The ability of a DNA to deform its structural characteristics under a specific stimulus is known as DNA bendability. A tri-nucleotide model based on DNAse-I cutting frequencies predicts that DNAse I binds and cuts DNA that is bent towards major groove [13, 15].

B. Thermodynamic signals: Stacking energy

The stability of both of a given DNA sequence and as well as protein interactions is indicated by the stacking energies which is also responsible for formation of local structures [13, 16].

C. Duplex stability: Free energy signals

The duplex stability of a DNA structure depends upon its base sequence and more specifically upon ten different types of nearest neighbour interactions namely AA/TT; AT/TA; CA/GT; GT/CA; CT/GA; GA/CT; CG/GC; GC/CG; GG/CC^13,14^

D. Propeller twist signals, Bending stiffness and Nucleosomal positioning

DNA must distort in order to bend around a protein: this distortion is facilitated by the deformational capacity of dinucleotides [8]. Some signals are rigid whereas others are flexible, and the propeller twist signals are responsible for measuring the flexibility of DNA [13, 17].

E. Protein interaction signals

The DNA sequences during their interaction with other molecules carries signals specific for its ability to deform [8].

### Results

Over 43.59% of gibbon genome is occupied by MGEs. The mobile genetic elements that were identified in the gibbon genome included the major transposable element (TE) classes that consist of long terminal repeat (LTRs), non-LTRs and DNA transposons. The figure 2 shows the distribution of transposable elements on different chromosomes of the gibbon genome. Non-LTR transposons were the most abundant TEs in the gibbon genome. Several varieties of super families of LINEs and SINEs were discovered in the gibbon genome. We find that SINEs were present in the form of two super-families namely, ALUs and MIRs, whereas LINE consisted of LINE1, LINE2 and LINE3/CR1. Among the LINEs, the L1s covered the major proportion of the gibbon genome, whereas in the case of SINEs, the ALUs were found to be in majority.

**Figure 2:**
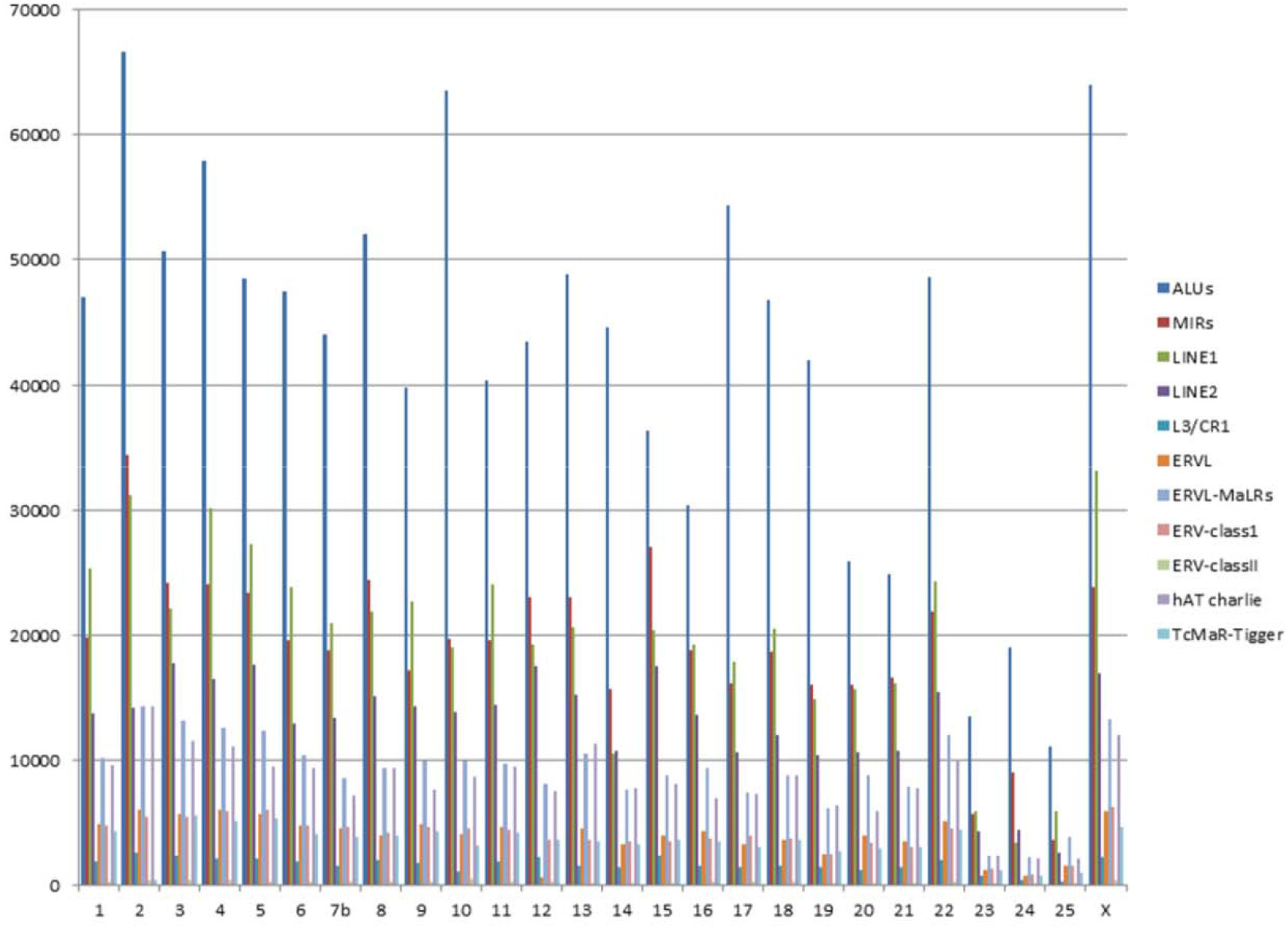
The chromosome wise representation of the distribution of MGEs in the gibbon genome. Different classes of elements are indicated with different colours. The y-axis represents the frequency of elements found on different chromosomes.

LTR retrotransposons were the second most abundant TEs in the gibbon genome and the majority of these belong to a class of mammalian repeats derived from retrovirus like elements. They were categorized into 4 subgroups namely; ERVL, ERVL-MaLRs, ERV-class I and ERV class II. DNA transposons are rare in the gibbon genome, and are represented by only two super-families (hAT-Charlie and TcMar-Tigger).

The table 1 shows the distribution of the mobile genetic elements in the different chromosomes of the gibbon genome. The percentage of each chromosome region occupied by repetitive elements was also calculated (Table 1). It was found out that ALUs were the most abundant transposable elements present in the gibbon whereas, LINE 1 was also one of the most frequently occurring MGE in the genome. Since there are large number of full length Alu copies in various genomes, this makes them suitable candidates for detailed analysis as representative SINEs.

**Table I:**
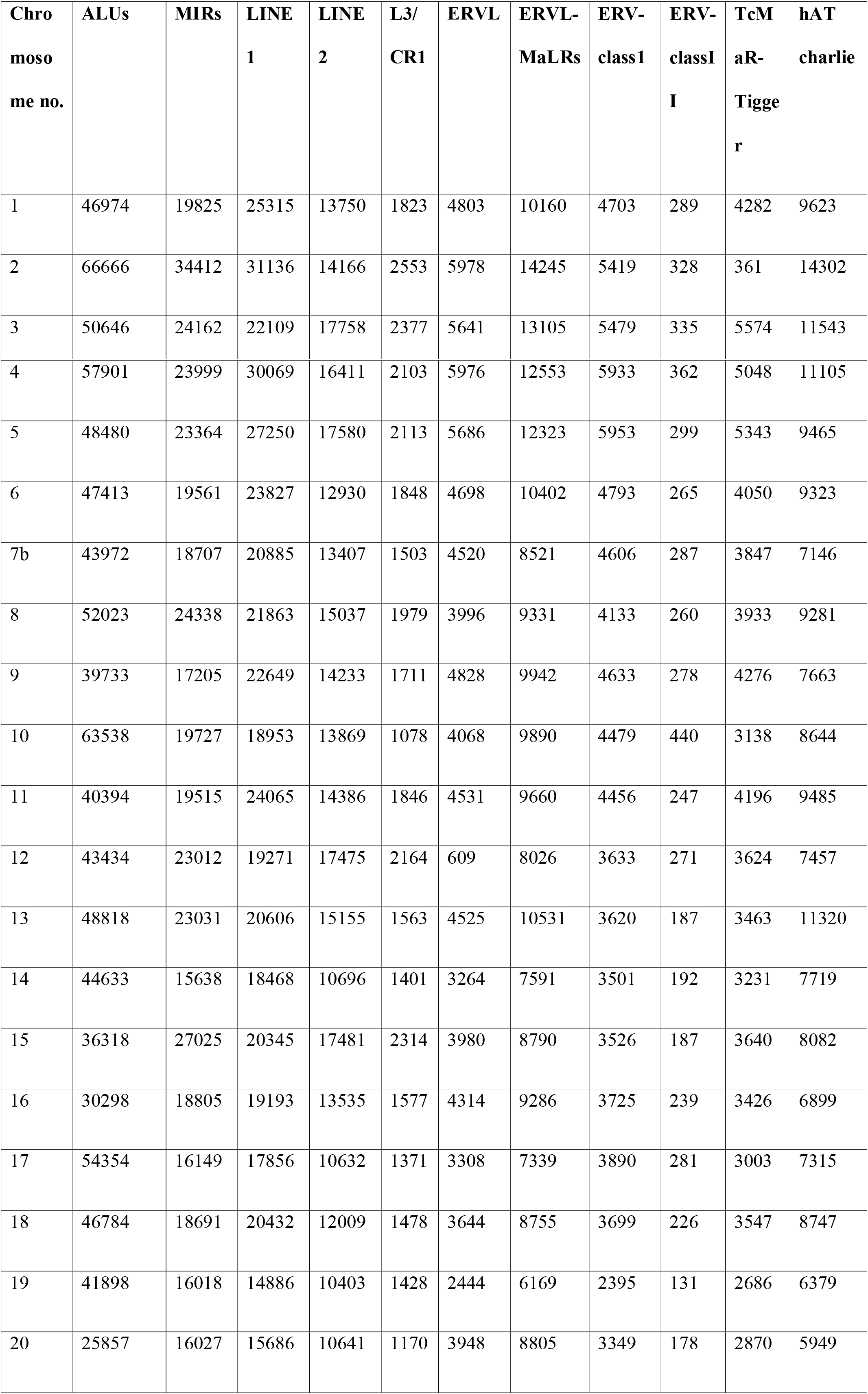
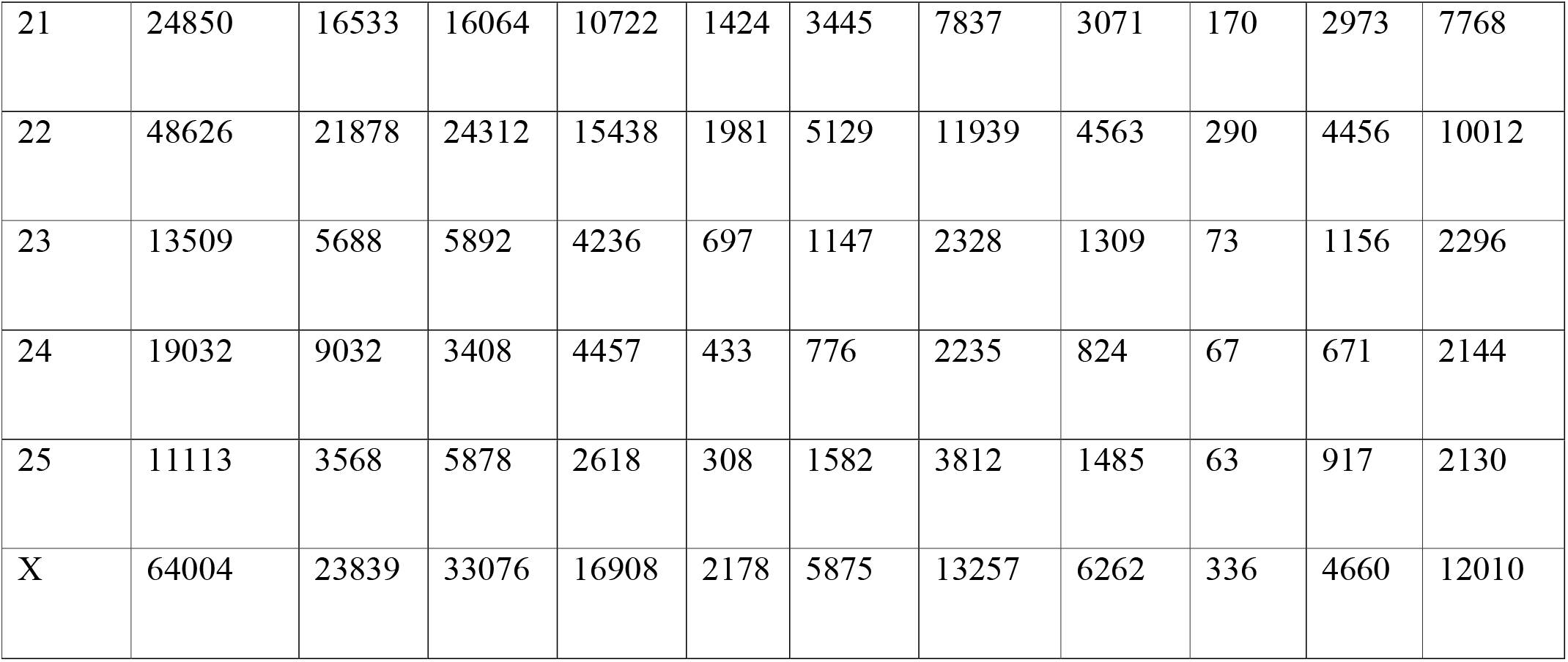
Distribution of transposable elements in the gibbon genome.

### Analysis of the LINEs and SINEs in gibbon genome

We performed detailed analysis of LINEs, SINEs and LTR elements for each chromosome using Perl scripts and results are summarised in Tables 2, 3 and 4. The total number of SINE elements were found to be 16, 07,203; LINE elements were 9, 31,127 in number and the total count of DNA elements were found to be 4, 12,435.

**Table II:** Chromosome wise summary of the SINE elements in gibbon genome.

| **Chromosome no.** | **Total length** | **Length occupied** | **No. of elements** | **Percentage of sequence (%)** |
| --- | --- | --- | --- | --- |
| 1 | 124228569 | 15389374 | 67140 | 12.39 |
| 2 | 164513156 | 22689809 | 101439 | 13.79 |
| 3 | 158281636 | 17116535 | 75264 | 10.81 |
| 4 | 154836434 | 18768321 | 82256 | 12.12 |
| 5 | 144113614 | 16384550 | 72266 | 11.36 |
| 6 | 122129856 | 15418127 | 67273 | 12.62 |
| 7b | 116018537 | 14380977 | 63042 | 12.30 |
| 8 | 117808995 | 17294411 | 76623 | 14.68 |
| 9 | 118737704 | 13153091 | 57266 | 11.07 |
| 10 | 108686836 | 19369927 | 83444 | 17.82 |
| 11 | 123853985 | 13649407 | 60277 | 11.02 |
| 12 | 109700369 | 14886328 | 66775 | 13.50 |
| 13 | 109844878 | 16187346 | 72089 | 14.7 |
| 14 | 99334225 | 14000626 | 60488 | 14.09 |
| 15 | 110683687 | 13749624 | 63638 | 12.42 |
| 16 | 100086279 | 10888927 | 49331 | 10.88 |
| 17 | 101490674 | 16425099 | 70723 | 16.18 |
| 18 | 105523347 | 9089854 | 65727 | 8.61 |
| 19 | 82726427 | 13371447 | 58100 | 16.16 |
| 20 | 85254692 | 9254553 | 42083 | 10.85 |
| 21 | 87080223 | 9095777 | 41643 | 10.44 |
| 22 | 133238221 | 16175660 | 70911 | 12.14 |
| 23 | 33724709 | 4909473 | 19334 | 14.55 |
| 24 | 28896242 | 6251379 | 28111 | 21.63 |
| 25 | 32264544 | 3484990 | 14759 | 10.80 |
| X | 144882752 | 18818075 | 82201 | 12.98 |

**Table III:** Chromosome wise summary of the LINEs in gibbon genome.

| **Chromosome no.** | **Total length** | **Length occupied** | **No. of elements** | **Percentage of sequence** |
| --- | --- | --- | --- | --- |
| 1 | 124228569 | 26587344 | 41596 | 21.4 |
| 2 | 164513156 | 32458804 | 55708 | 19.73 |
| 3 | 158281636 | 33196243 | 51486 | 20.97 |
| 4 | 154836434 | 29872691 | 49351 | 19.29 |
| 5 | 144113614 | 28093514 | 48012 | 19.4 |
| 6 | 122129856 | 23513619 | 38586 | 19.25 |
| 7b | 116018537 | 20850749 | 36404 | 17.97 |
| 8 | 117808995 | 21855513 | 39996 | 18.5 |
| 9 | 118737704 | 25798628 | 39250 | 21.72 |
| 10 | 868683610 | 17529436 | 34306 | 16.12 |
| 11 | 123853985 | 26695501 | 41001 | 21.55 |
| 12 | 109700369 | 23387137 | 39489 | 21.3 |
| 13 | 109844878 | 18859033 | 37758 | 17.16 |
| 14 | 99334225 | 17941765 | 31005 | 18.06 |
| 15 | 110683687 | 24119300 | 40681 | 21.79 |
| 16 | 100086279 | 21904114 | 34870 | 21.88 |
| 17 | 101490674 | 17468268 | 30361 | 17.21 |
| 18 | 105523347 | 20511590 | 34469 | 19.44 |
| 19 | 82726427 | 13787352 | 27121 | 16.67 |
| 20 | 85254692 | 16821821 | 27910 | 19.73 |
| 21 | 87080223 | 17954893 | 28634 | 20.62 |
| 22 | 133238221 | 25149791 | 42499 | 18.87 |
| 23 | 33724709 | 6844934 | 11032 | 20.2 |
| 24 | 28896242 | 3087909 | 8407 | 10.69 |
| 25 | 32264544 | 5486671 | 8953 | 17 |
| X | 144882752 | 33881737 | 52842 | 23.38 |

**Table IV:** Chromosome wise summary of the DNA elements in gibbon genome.

| Chromosome no. | Total length (bp) | Length occupied | No. of elements | Percentage of sequence |
| --- | --- | --- | --- | --- |
| 1a | 124228569 | 4578519 | 18734 | 3.69 |
| 2 | 164513156 | 5981178 | 25705 | 3.64 |
| 3 | 158281636 | 5799716 | 23205 | 3.67 |
| 4 | 154836434 | 5324876 | 21768 | 3.43 |
| 5 | 144113614 | 5116564 | 20062 | 3.55 |
| 6 | 122129856 | 4268588 | 18098 | 3.49 |
| 7b | 116018537 | 38518008 | 14800 | 3.31 |
| 8 | 117808995 | 4136569 | 17769 | 3.51 |
| 9 | 118737704 | 4192221 | 16048 | 3.53 |
| 10 | 868683610 | 3441747 | 15483 | 3.16 |
| 11 | 123853985 | 4518862 | 18372 | 3.64 |
| 12 | 109700369 | 3704003 | 15278 | 3.37 |
| 13 | 109844878 | 4361802 | 19462 | 3.97 |
| 14 | 99334225 | 3396239 | 14670 | 3.42 |
| 15 | 110683687 | 3863138 | 15987 | 3.49 |
| 16 | 100086279 | 3567688 | 14056 | 3.56 |
| 17 | 101490674 | 3106632 | 13797 | 3.06 |
| 18 | 105523347 | 3789619 | 16231 | 3.59 |
| 19 | 82726427 | 2818295 | 12191 | 3.41 |
| 20 | 85254692 | 2978639 | 12066 | 3.40 |
| 21 | 87080223 | 3307062 | 14011 | 3.79 |
| 22 | 133238221 | 4821418 | 19763 | 3.62 |
| 23 | 33724709 | 1158301 | 4730 | 3.43 |
| 24 | 28896242 | 751431 | 3739 | 2.60 |
| 25 | 32264544 | 1025009 | 4208 | 3.17 |
| X | 144882752 | 5092937 | 22202 | 3.51 |

### Analysis of LAVA element in gibbon genome

For each chromosome, the analysis of the LAVA was performed (Figures 8-12). It was observed that the signals present in the upstream region were not distinct as observed previously [6, 18, 17, 11]. The insertion sites of LAVA also did not show any characteristic pattern in comparison with signals observed at insertion sites of Alus. The reasons could be attributed to absence of full length LAVA elements, absence of 5’ intact copies or possibly distinct insertion mechanism in comparison to LINEs or SINEs.

### Identification of transposable elements using ELAN

We present insertion sites of Alus in chromosome 24 as a representative case. Signals upstream of the insertion sites of Alu in chromosome 24 were obtained from DNA SCANNER. It has been observed that Alu elements tend to insert preferentially in the A-rich regions where parameters like thermodynamic properties, structural features and nucleosomal positioning parameters are different as compared to genomic average values. In contrast, LAVA elements seem to show a different pattern at insertion sites.

The peaks or troughs were obtained using DNA scanner which compute physiochemical properties using sliding window mechanism. Different physiochemical properties like DNA bendability, bending stiffness, protein interaction signals, and frequency of AT etc. were computed. The y axis in the profile represents value of the property and the x-axis gives the relative position with respect to the insertion site (See Figures 3-12).

**Figure 3:**
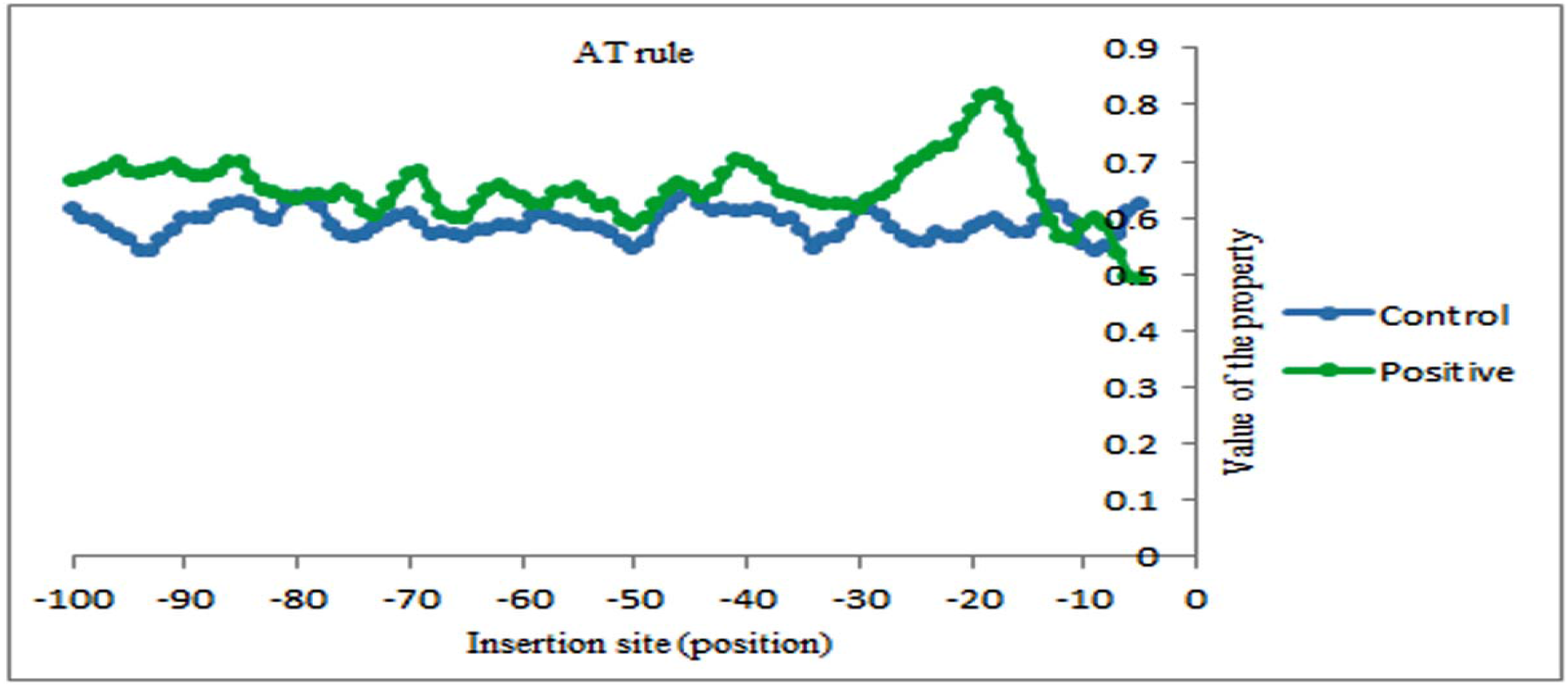
Adenine thymine density profile for the insertion sites present upstream of full length Alu insertions on 24^th^ chromosome of gibbon computed by DNA SCANNER. The y-axis represents the value of the property under study (referred as ‘AT Rule’) and the x-axis represents the position with respect to insertion site (taken as position 0). Here, profile plotted in blue represent control dataset (randomly selected genomic sites) and green curve shows profile for the positive dataset.

**Figure 4:**
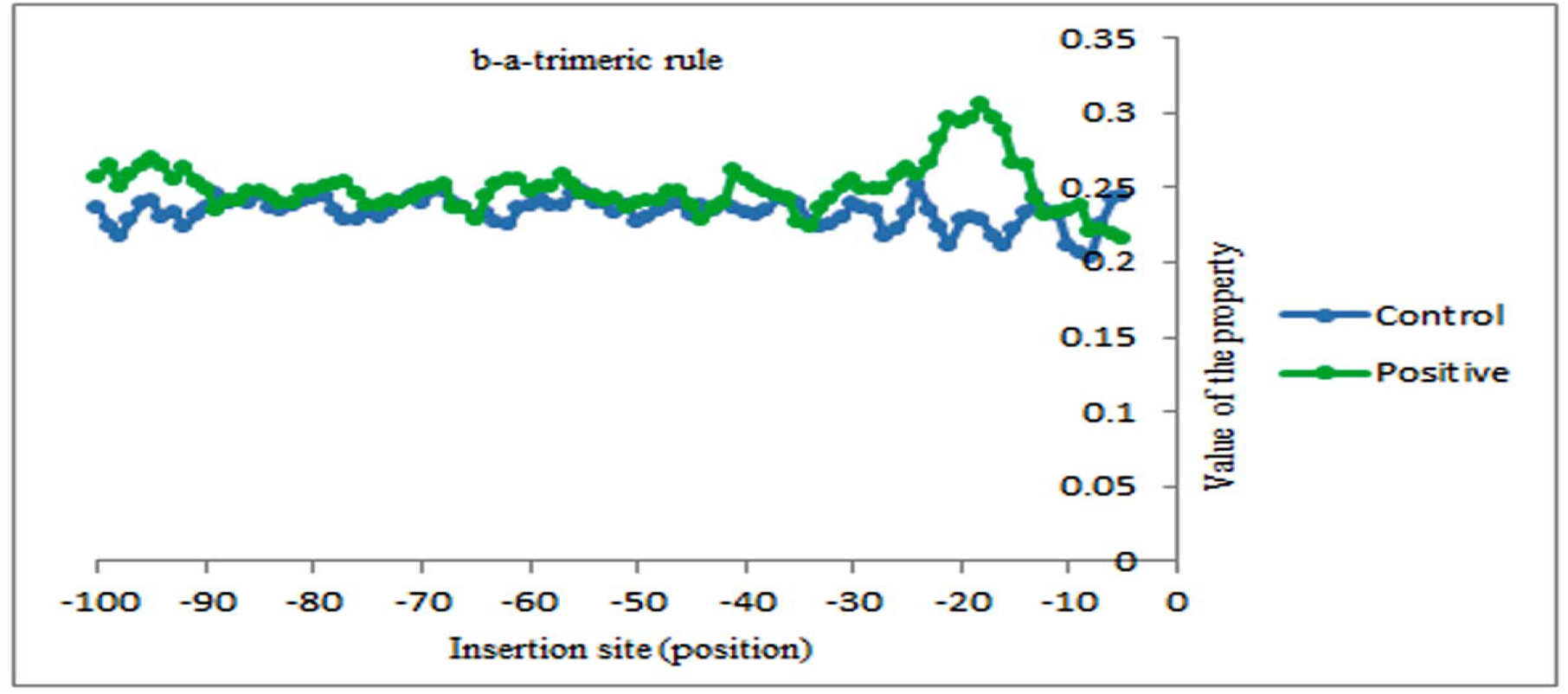
B to A form density parameter profile for the insertion sites present upstream of full length Alu insertions in chromosome 24 of gibbon genome.

**Figure 5:**
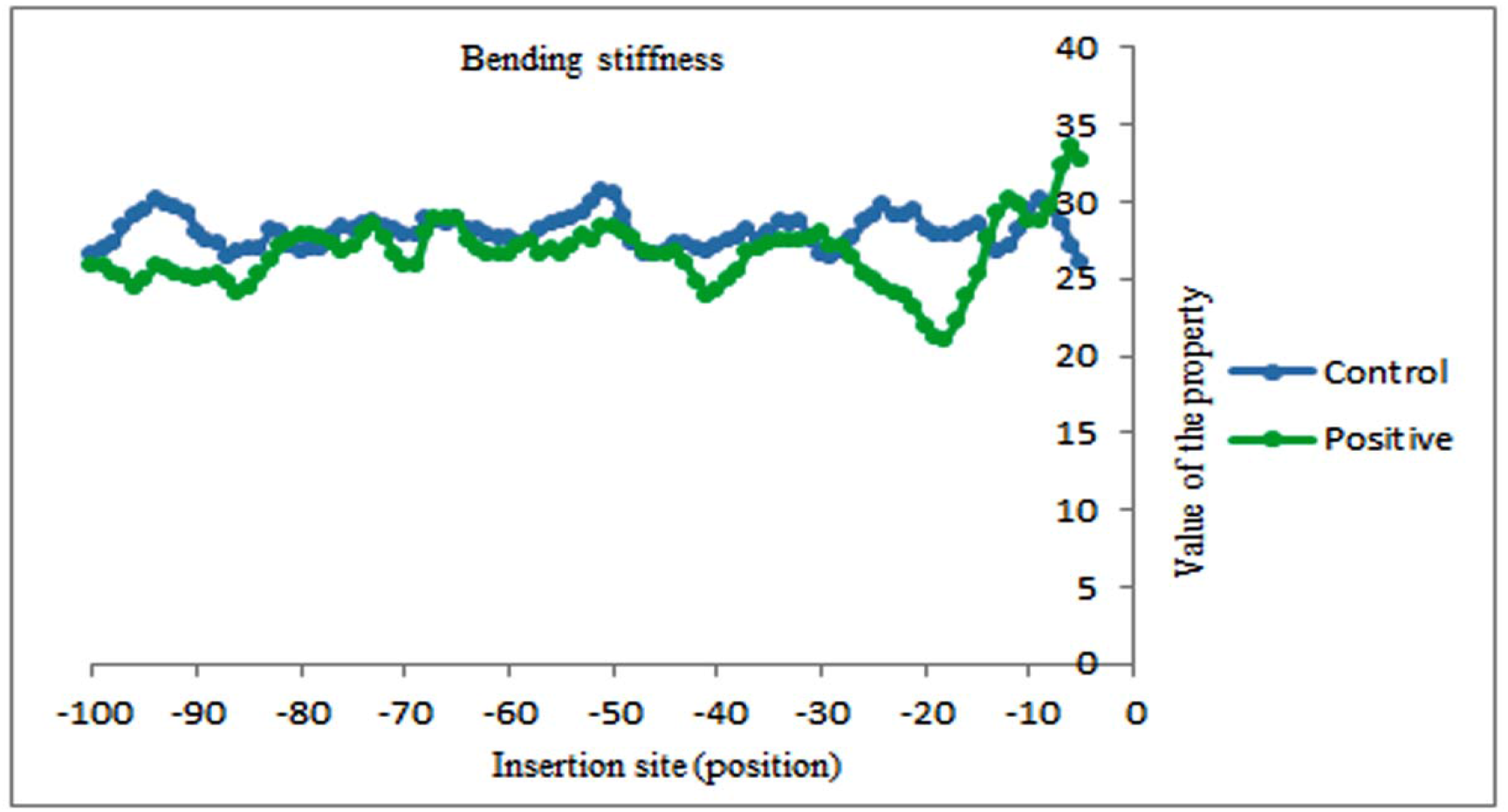
Bending stiffness profile for the insertion sites present upstream of full length Alu insertions in chromosome 24 of gibbon genome.

**Figure 6:**
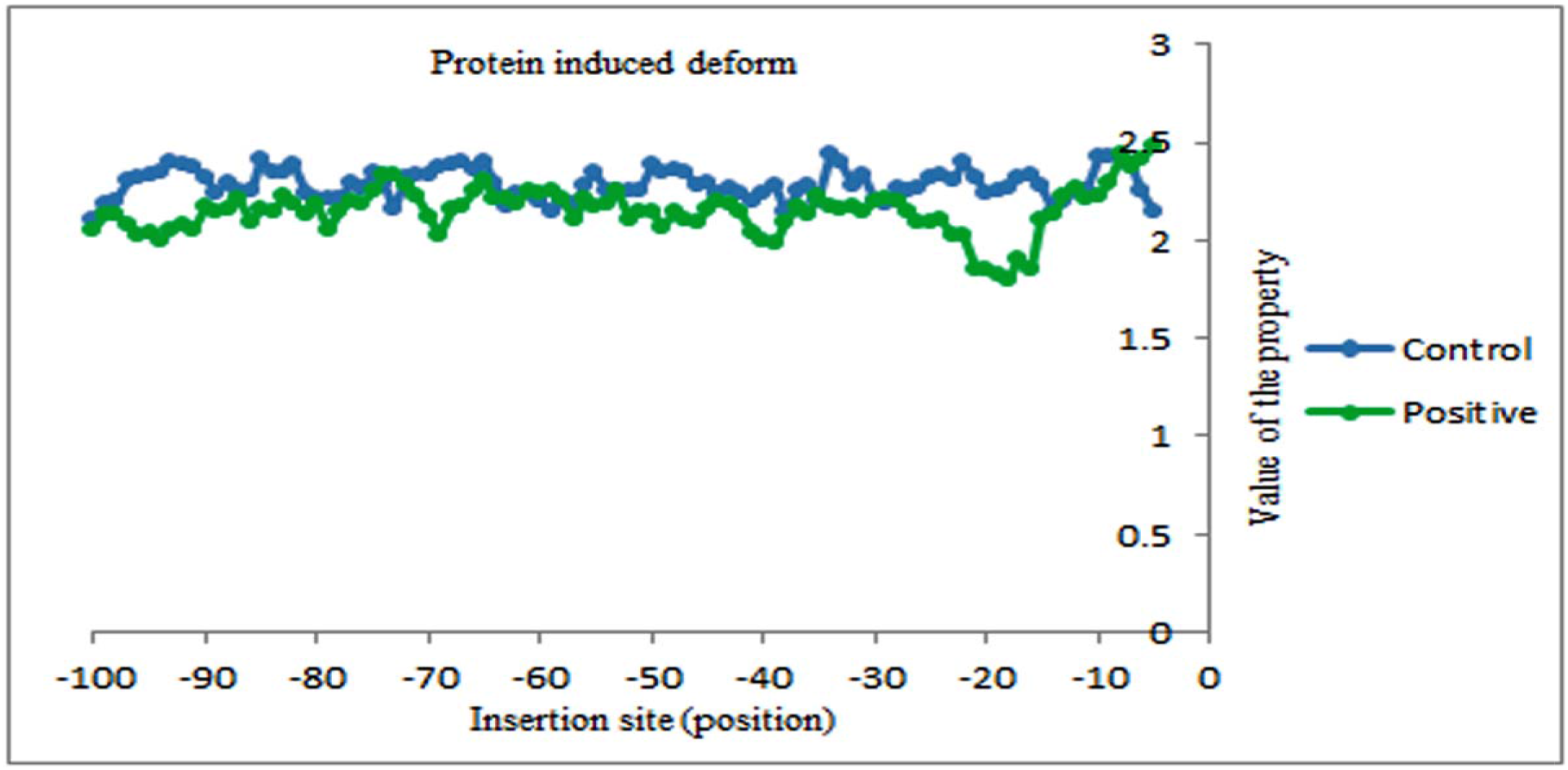
Protein induced deformability profile generated for the insertion sites present upstream of full length Alu insertions in chromosome 24 of gibbon genome.

**Figure 7:**
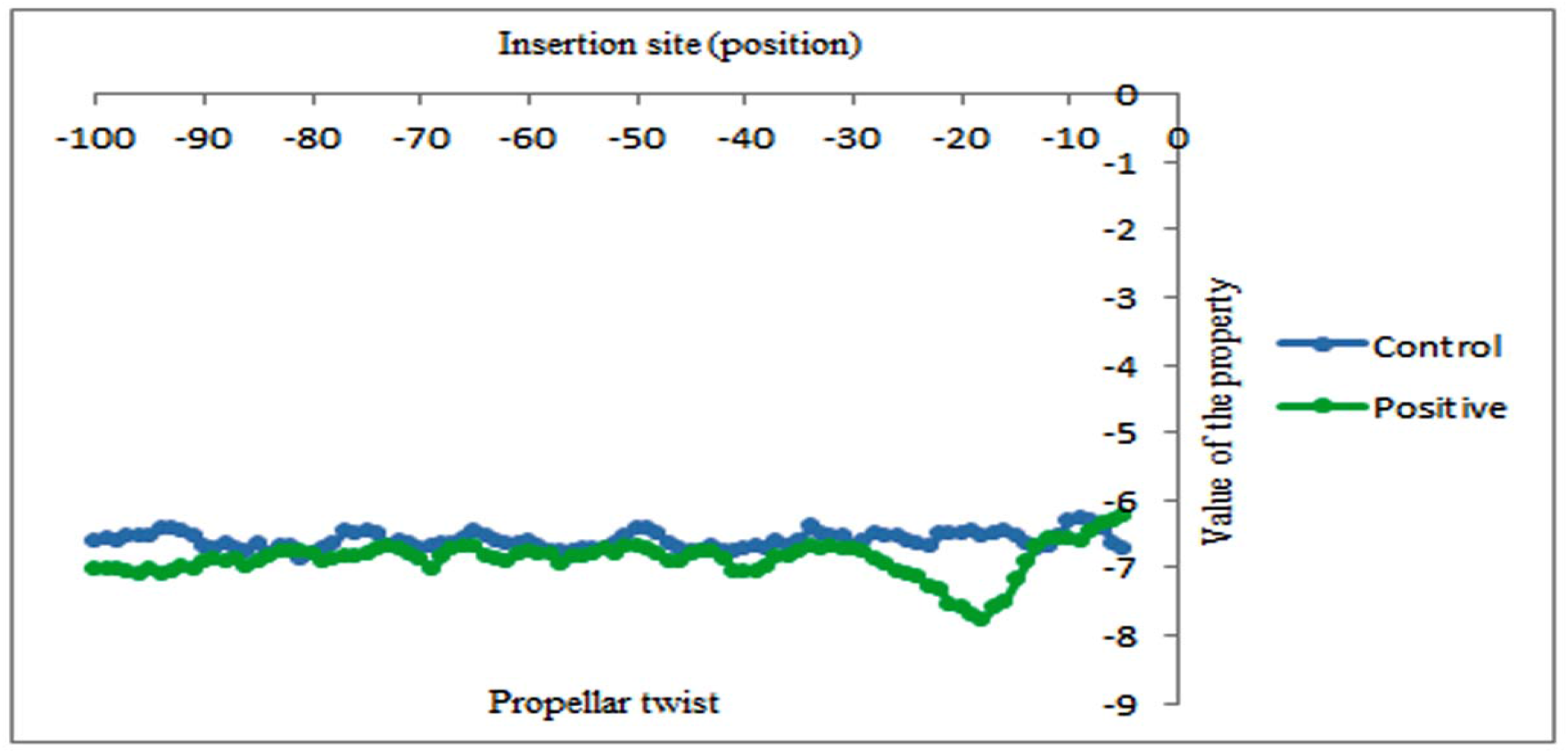
Propeller twist profile for the insertion sites present upstream of full length Alu insertions in chromosome 24 of gibbon genome.

**Figure 8:**
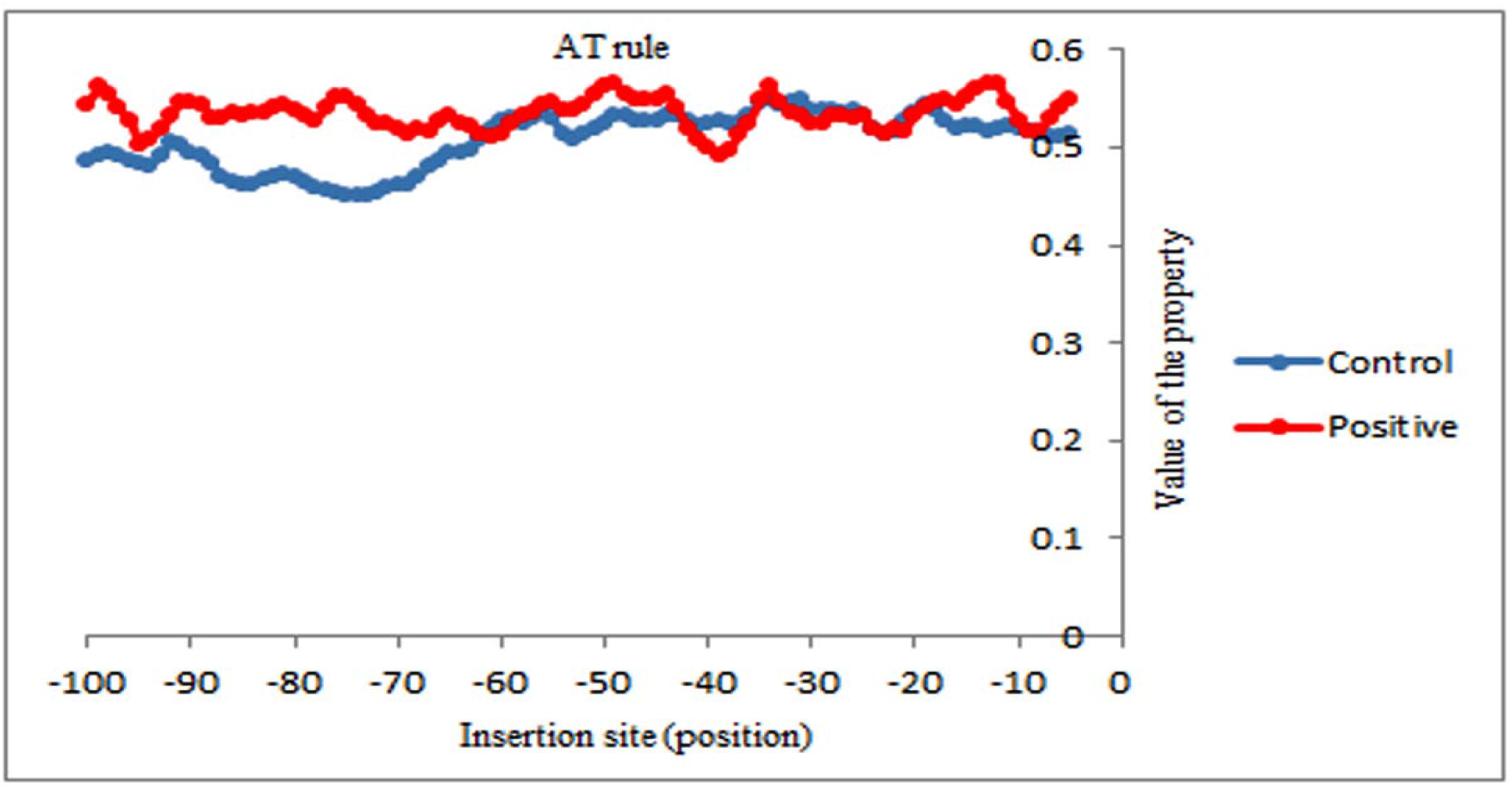
AT frequency profile for upstream region of LAVA insertions in chromosome 24.

**Figure 9:**
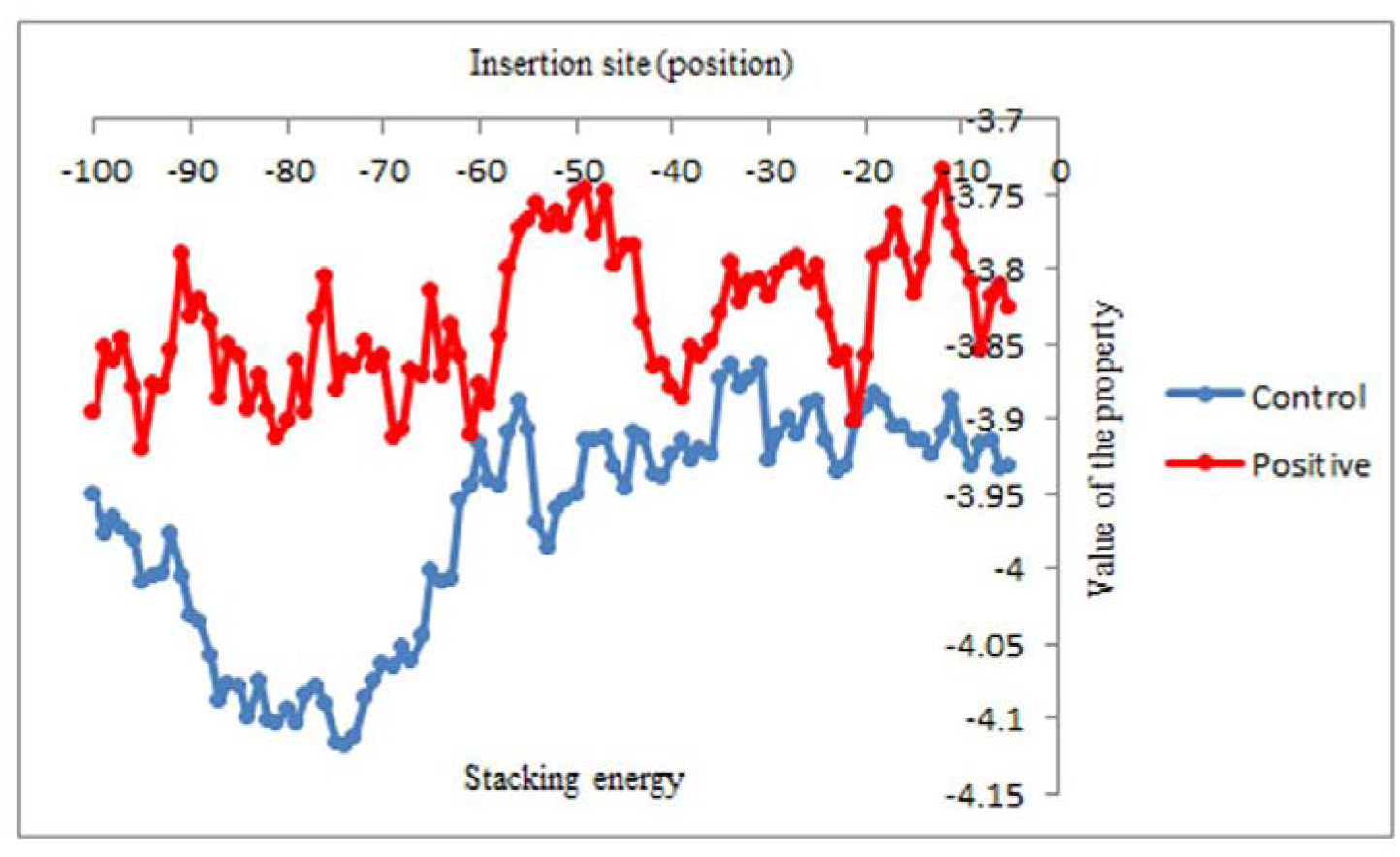
Stacking energy profile for upstream region of LAVA insertions in chromosome 24.

**Figure 10:**
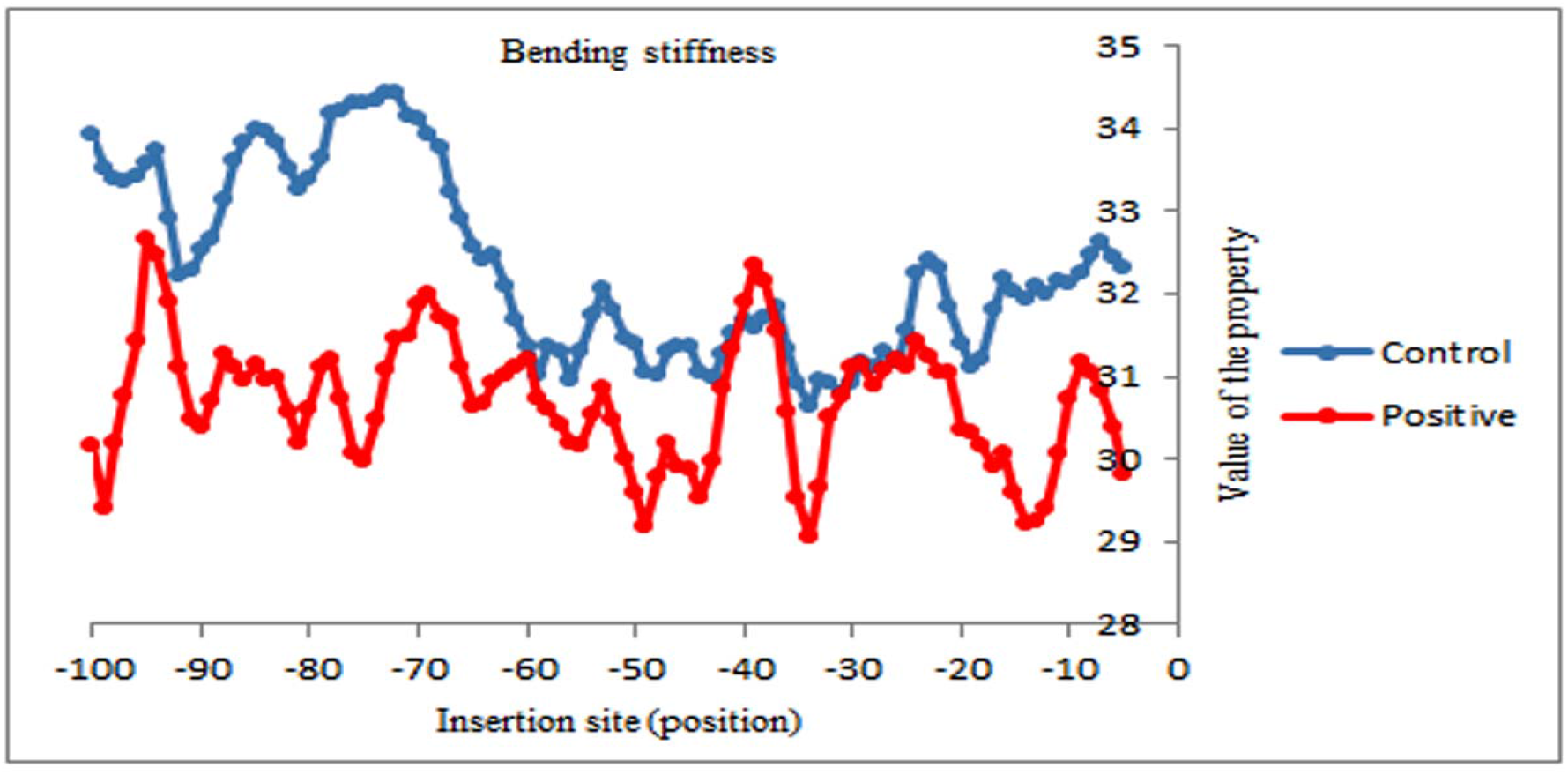
Bending stiffness profile for upstream region of LAVA insertions in chromosome 24.

**Figure 11:**
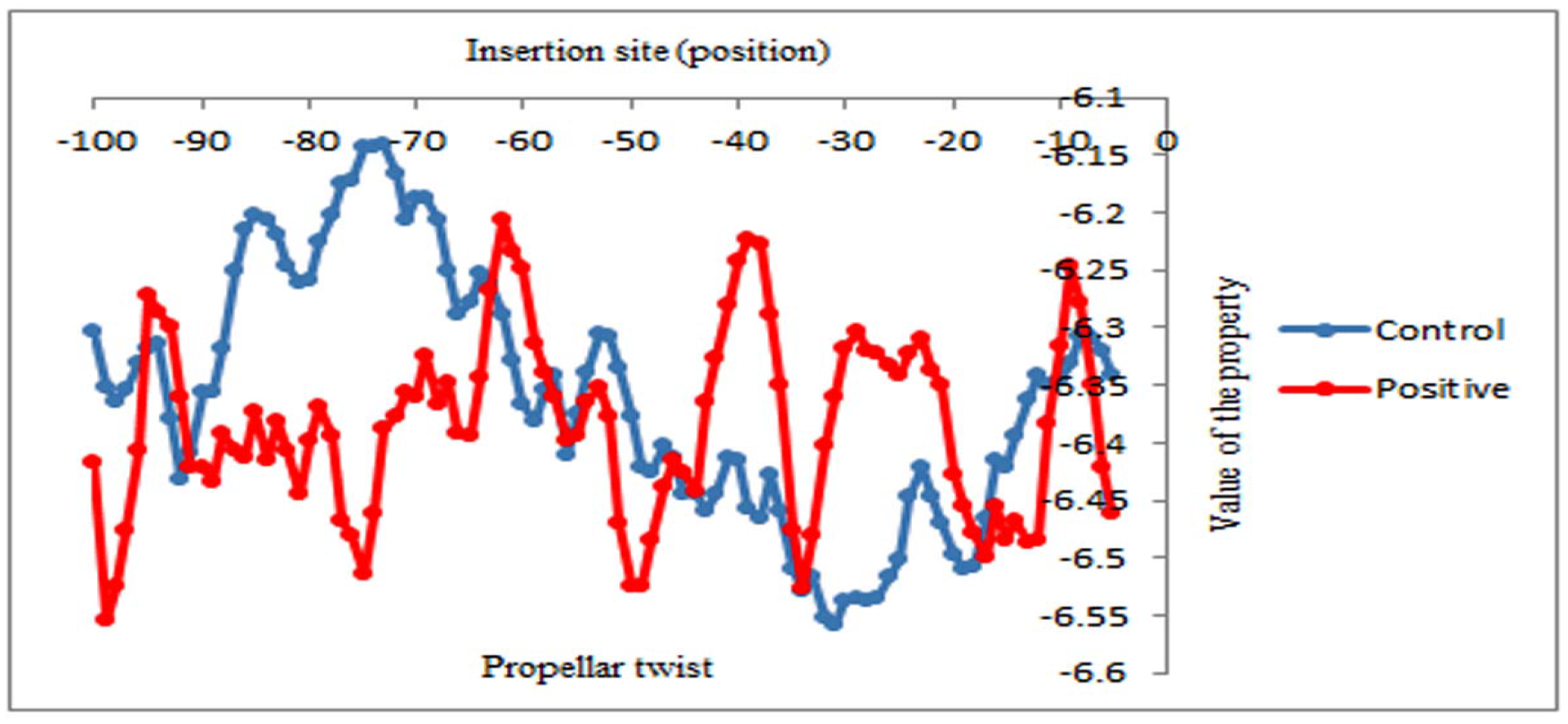
Propeller twist profile for upstream region of LAVA insertions in chromosome 24.

**Figure 12:**
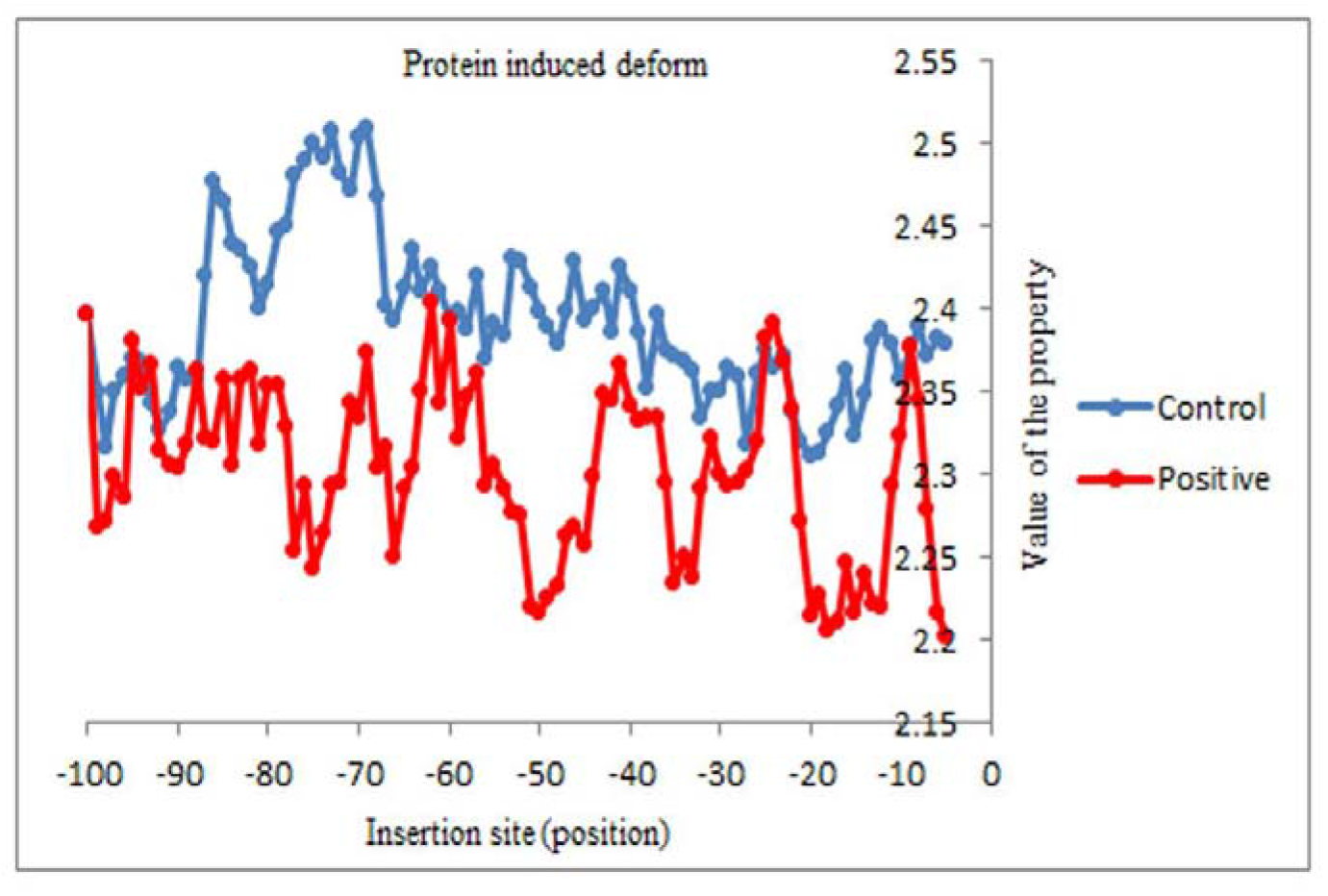
Protein induced deformability profile for upstream region of LAVA insertions in chromosome 24.

### Discussion

Recent sequencing of gibbon has contributed significantly in understanding evolution of human, chimpanzee and gorilla. The gibbon genome contains all previously described classes of transposable elements that are present in other primates as well. The mystery surrounding transposable elements revolves around their mode and selection of insertion site. Different categories of transposable elements uses different mechanism to get inserted into the genome which includes target prime retrotransposition (TPRT). Previously, gibbon-great ape population divergence time was estimated to be of ~16.8 million years ago assuming a split time with macaque of 29 million years ago, therefore the LAVA element is believed to be originated around the time of the divergence of gibbons from the ancestral great ape or human lineage. The LAVA element is implicated for mechanism to cause gross chromosomal changes leading to accelerated evolution.

We have used Repeatmasker and ELAN to find different mobile genetic elements from gibbon genome. In ELAN, ELEFINDER and DNASCANNER were used for detailed analysis of insertions of Alu and LAVA elements in the genome and to scan the insertion sites of gibbon chromosomes for different physicochemical properties for protein interactions [8]. We observed that there is a significant difference in patterns or signals observed in insertion sites of Alus and LAVA elements. This suggests that LAVA element tend to insert in a mechanism which is distinct from the mechanism and property of LINEs and SINEs. Previously, we have shown that signals observed in insertion sites of Alu elements are different when inserted in exonic region, intronic as well as intergenic regions [19]. The signals observed in intergenic regions are conserved across various Alu insertion sites in various primate genomes [13, 19, 20]. Previously, we conducted detailed analyses of genes in the research which included genes like APC, DMD and CYBB for Alu, L1 and SVA insertions that were reported to be responsive to TE insertion leading to disease. In our study, we found that the signals observed in most of the disease genes were atypical in nature like in DMD which is one of the largest genes known of 2.4 Mb in length. Also, it was seen that there is an occurrence of L1 insertion which has been reported to cause disruption in the gene which further leads to Duchene myotrophy disease. This and other examples can be seen online at *http://nldsps.jnu.ac.in/TE-Vs-diseases/database/*. As per our study of 49 genes, we assume that the absence of typical signals within exonic regions prevents further wide-spread disruption of coding regions by Alu and L1s respectively. Similarly, it was suggested by Carbone et al 2014 that the high level of chromosomal rearrangements in the gibbon genome may have been due to the premature transcription termination mechanism of the segregated genes of the chromosomes due to LAVA activities. Though this mechanism occurred at a very low level but still it was sufficient to increase the chromosome segregation errors [9]. Hence, it is interesting to observe that the possible genic targets sites of LAVA element presents atypical signals that are quite similar to the signals observed in human genes amenable for disease causing MGE insertions. . Importantly, these signals were conserved in genomes of the lower eukaryotes such as *E. histolytica* [20].

## Acknowledgement

We acknowledge contribution of our lab members during the preparation of this manuscript: Hitesh Jaiswal, Samridhi Mehta ^1*^, Garima Chaudhary ^1^, Mitika Raj ^1^, Srashti Varshney ^1^, Shubhani Chokra. We also acknowledge DBT, Government of India for their support. We also thanks Mr. Vishal Kumar Mishra for the help extended during the submission process. We also thank anonymous reviewers for their valuable comments.

